# Modeling the Impact of Dynamic Gastric pH on *Helicobacter pylori* Eradication and Antibiotic Resistance Emergence

**DOI:** 10.64898/2026.02.26.708110

**Authors:** Alex Hermann Sockeng Koussok, Edward Richard Onyango, Koichi Fujimoto, Jean Jules Tewa

**Affiliations:** Department of Mathematics, Pan African University Institute for Basic Sciences, Technology and Innovation, Nairobi, 00200, Kenya; Department of Pure and Applied Mathematics, Jomo Kenyatta University of Agriculture and Technology, Nairobi, 00200, Kenya; Graduate School of Integrated Sciences for Life, Hiroshima University, Higashi-Hiroshima, 739-8528, Hiroshima, Japan; National Advanced School of Engineering of Yaounde, University of Yaoundé I, Yaoundé, P.O. Box 8390, Cameroon

**Keywords:** *H. pylori*, Within-host model, Antibiotic resistance, Gastric pH, Urease

## Abstract

*Helicobacter pylori* infections present a persistent global health challenge due to increasing antibiotic resistance and the bacterium’s ability to survive in the acidic gastric environment. Existing within-host models of H. pylori infection neglect the gastric pH fluctuation, despite its role in modulating bacterial growth and antibiotic efficacy. To address this gap, we extend a published in-host model by explicitly incorporating gastric pH as a dynamic state variable, influenced by three key physiological processes (i) bacteria urease which neutralizes gastric acid to create a protective niche; (ii) host acid secretion response, which attempts to restore baseline acidity; and (iii) dietary perturbations, which induce temporary pH changes. Equilibrium and stability analysis reveal pH-dependent reproductive thresholds ℛ_***s***_**(*H*)** and ℛ_***r***_**(*H*)** that determine the conditions for bacterial persistence and treatment outcome. Successful eradication requires driving both thresholds below unity. Numerical simulations validate distinct clinical scenarios including complete bacterial clearance, resistant strain dominance, stable bacterial coexistence, and oscillatory persistence. These outcomes emerge from the coupled interplay between antibiotic pressure, immune response, and pH regulation. Our model provides a comprehensive theoretical framework for understanding *H. pylori* treatment failures and highlights how adjuvant pH-modulation strategies could enhance antibiotic efficacy against resistant infections.

## 1 Introduction

*Helicobacter pylori*, a gram-negative bacterium that colonizes the human gastric mucosa, is a major etiological agent of chronic gastritis, peptic ulcer disease, and even gastric cancer [1, 2]. Its persistence in the harsh, acidic environment of the stomach is a remarkable feat of microbial adaptation, primarily mediated by the enzyme urease which locally neutralizes gastric acid, creating a survivable niche [3]. Standard eradication therapy typically involves a combination of a proton pump inhibitor and two antibiotics, yet treatment failure rates remain alarmingly high, ranging from 20% to 40% globally [4]. This therapeutic challenge is compounded by the alarming rise in antibiotic resistance to first-line agents like clarithromycin and metronidazole [5].

The complex dynamics of *H. pylori* infection and treatment failure operate across multiple scales, from the molecular mechanisms of antibiotic resistance to the ecological interactions within the gastric environment. Within-host mathematical models provide a powerful framework to synthesize these mechanisms and explore their integrated effects on treatment outcomes [6]. Existing models have successfully captured key features such as the dynamics of colonization in relation to the host response [7], the competition between sensitive and resistant bacterial populations under antibiotics therapy [8], and a recent extension with modulating role of the host immune response [9].However, these models did not consider the explicit dynamic role of gastric pH.

Gastric pH is not a static, hostile barrier but a dynamically regulated variable that is actively manipulated by both the host and the pathogen. *H. pylori*’s urease activity elevates the local pH, facilitating colonization [10]. In response, the host’s acid-secretory machinery attempts to restore homeostasis [11]. Furthermore, exogenous factors like meals and acid-suppressive drugs (PPI) introduce time-dependent pH fluctuations. This dynamic pH environment profoundly influences bacterial growth rates [12], the chemical stability and efficacy of antibiotics [13, 14], and potentially the activity of immune cells. Ignoring this critical axis limits the predictive power and clinical relevance of within-host models, as it omits a primary mechanism by which bacteria evade both host defenses and antimicrobial therapy.

To bridge this gap, we extend the within-host model developed by Noguera et al. [9], which elegantly describes the competitive dynamics of sensitive and resistant *H. pylori* strains under antibiotic pressure with an adaptive immune response. Our contribution is the explicit incorporation of gastric pH (*H*(*t*)) as a dynamic state variable. The extended model mechanistically couples: (i) bacterial urease activity, raising pH proportional to the total bacterial load; (ii) host homeostatic acid secretion, modeled as negative feedback towards a baseline acidic pH; and (iii) dietary perturbations as time-dependent inputs. The bacterial growth rates (*β*_*S*_, *β*_*R*_) and antibiotic killing efficacies (*κ*_*i*_) are therefore functions of pH, based on established experimental data [12, 14].

We perform a complete analytical and numerical investigation of this pH-integrated model. We derive non-dimensionalized equations and identify all biologically feasible equilibria: the infection-free state, resistant-only states (with and without immunity), and coexistence states. Stability analysis yields pH-dependent basic reproductive numbers, ℛ _*s*_(*H*) and ℛ _*r*_(*H*), which serve as critical thresholds for bacterial persistence. We demonstrate that successful eradication requires driving both thresholds below unity, a condition that depends intricately on the coupled system’s state. Through comprehensive numerical simulations, we map parameter spaces to distinct clinical outcomes successful clearance, resistant strain dominance, stable bacterial coexistence, and oscillatory persistence revealing how the interplay between antibiotic action, immune clearance, and pH modulation dictates treatment fate.

This work provides a more physiologically grounded and comprehensive theoretical framework for understanding *H. pylori* treatment outcomes. It highlights gastric pH not merely as a background condition but as a central, dynamic player in the host-pathogen-therapy triad. Our findings underline the rationale for acid-suppressive therapy in eradication regimens and suggest that precise manipulation of the gastric pH landscape could be a strategic target for improving antibiotic efficacy, particularly against resistant infections. By integrating this key ecological variable, the model offers a refined tool for exploring and optimizing therapeutic strategies against this pervasive and resilient pathogen.

## 2 Model Formulation

In this section, we extend the within-host model of *H. pylori* infection by Noguera et al. [9] to incorporate dynamic gastric pH as a critical ecological determinant of bacterial persistence and antibiotic efficacy. The based framework described interactions between sensitive (*S*(*t*)) and resistant (*R*(*t*)) bacterial populations under antibiotic therapy and immune response (*T* (*t*)). Our extended model introduces gastric pH (*H*(*t*)) as a dynamic variable influenced by three key physiological processes: bacterial urease activity, host acid-secretion feedback [11], and dietary perturbations.

The bacterial populations *S*(*t*) and *R*(*t*) represent densities of antibiotic-sensitive and antibiotic-resistant *H. pylori*, respectively. The immune response *T* (*t*) models the population of *T* lymphocytes involved in bacterial clearance. Three antibiotic concentrations *D*_*i*_(*t*) (for *i* = 1, 2, 3, representing clarithromycin, rifampicin, and Ciprofloxacin) follow standard pharmacokinetics. The novel component, gastric pH *H*(*t*), is modeled as a continuous variable ranging from the baseline acidic state (*H*_0_ ∈ [1.5, 3.5]) to the maximum alkalinization achievable through bacterial urease activity (*H*_max_ ≈ 7) [3, 15].

The complete dynamics of growth and *H. pylori* resistance acquisition as well as the immune response is represented by Figure 1 with the gastric pH as a dynamic variable (*H*(*t*)) that was not present in previous frameworks. Solid arrows represent fluxes between compartments, while self-loops indicate intrinsic growth or dynamics.

**Fig. 1.**
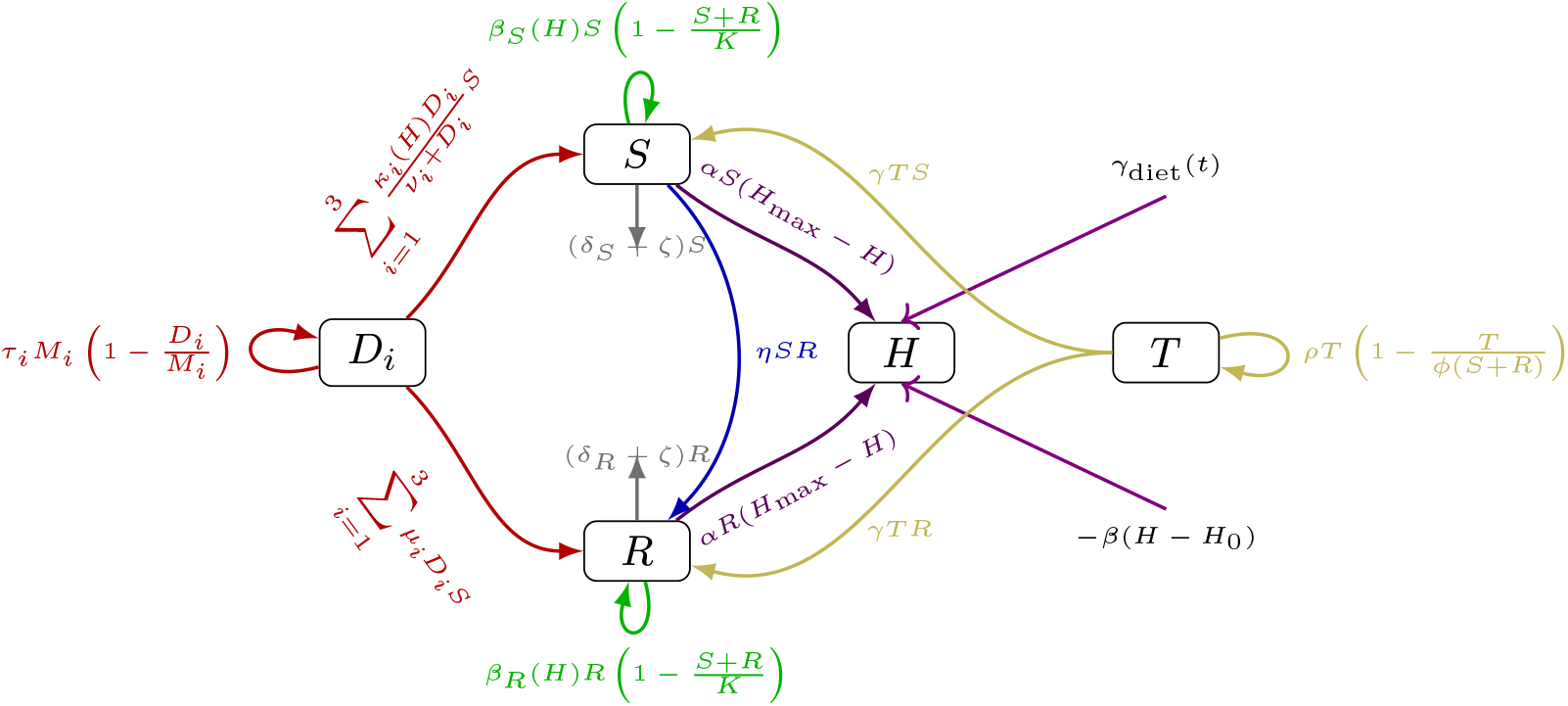
Schematic representation *H. pylori* infection model under antibiotic therapy, with immune response and dynamic pH regulation. Colors denote different biological processes: green (bacterial growth), red (antibiotic effects), blue (plasmid transfer), yellow (immune response), violet (pH modulation), and gray (bacterial death/detachment).

The dynamics described by Figure 1 are governed by the following system of nonlinear ordinary differential equations:

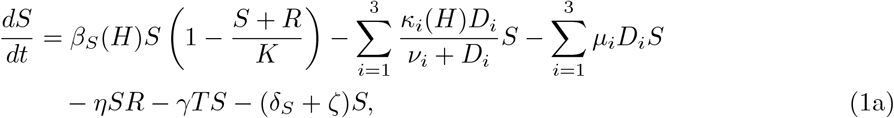

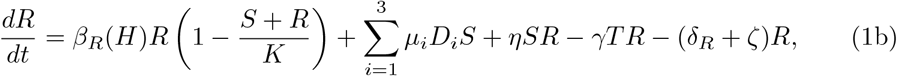

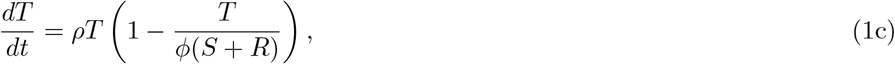

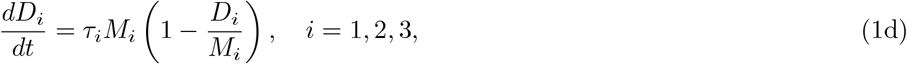

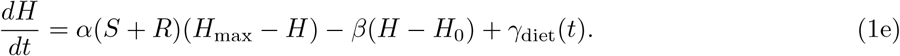

Bacterial population growth is logistic with a shared capacity *K*. The growth rates *β*_*S*_(*H*) and *β*_*R*_(*H*) are explicit functions of pH, capturing the known pH optimum of *H. pylori* growth around neutral pH [12] and the symmetric decrease when the pH deviates from neutrality. Antibiotic action against sensitive bacteria follows saturable Michaelis-Menten kinetics with maximum killing rates *κ*_*i*_(*H*) that also depend on pH, reflecting pH-dependent antibiotic stability and activity [14]. The terms *µ*_*i*_*D*_*i*_*S* represent mutation from sensitive to resistant phenotypes under antibiotic pressure, while *ηSR* models horizontal gene transfer. Immune-mediated clearance occurs at rate *γ*, and lymphocyte proliferation follows a density-dependent recruitment term limited by the total bacterial load.

The novel pH dynamics in equation (1e) incorporate three physiological mechanisms:

1. bacterial urease activity proportional to total bacterial density (*S* + *R*), which alkalinizes the environment toward *H*_max_ at rate *α*; under the assumption that both sensitive and resistant strength contribute equally to acid neutralization.
2. host compensatory acid secretion that restores baseline pH *H*_0_ at rate *β*; assumed following a negative linear feedback around *H*_0_.
3. time-dependent dietary perturbations *γ*_diet_(*t*) representing meal-induced pH changes assume to be time varying and decaying exponentially. The dietary term follows:

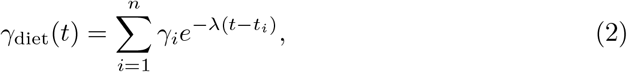

where each meal at time *t*_*i*_ introduces an instantaneous pH change of magnitude *γ*_*i*_ that decays exponentially with rate *λ*.

A key innovation is the explicit pH-dependence of bacterial growth and antibiotic efficacy. Following experimental data [12], bacterial growth rates follow:

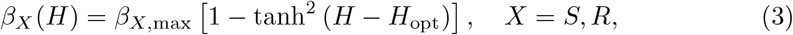

with optimal pH *H*_opt_ = 7.0. Antibiotic efficacy exhibits Gaussian dependence on pH:

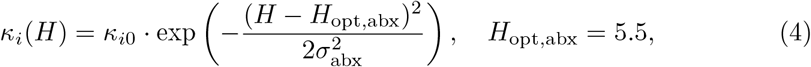

reflecting maximal antibiotic activity in the moderately acidic range achieved during proton pump inhibitor therapy [16].

To facilitate the analysis, we non-dimensionalize the system using scale factors: bacterial populations by carrying capacity *K*, immune cells by *ϕK*, antibiotics by their maximum concentrations *M*_*i*_, and pH relative to its dynamic range. From the change of variables *s* = *S/K, r* = *R/K, g* = *T/*(*ϕK*), *d*_*i*_ = *D*_*i*_*/M*_*i*_, and *h* = (*H* − *H*_0_)*/*(*H*_max_ − *H*_0_), we obtain the reduced system:

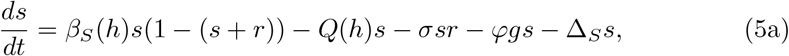

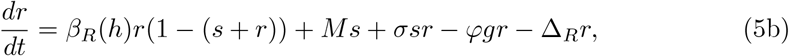

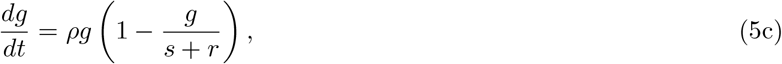

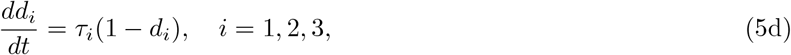

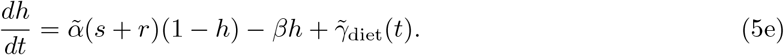

The composite parameters are defined as:

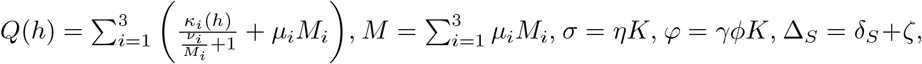

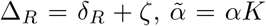, and 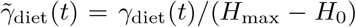. The system evolves in the biologically relevant domain

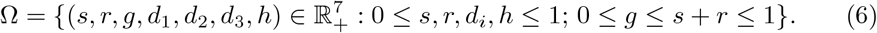

This pH-extended model captures the essential feedbacks between bacterial growth, antibiotic action, immune response, and gastric acidity that determine infection outcomes. The inclusion of pH dynamics allows investigation of how acid-suppressive therapies, dietary patterns, and bacterial urease activity jointly influence treatment efficacy and resistance emergence.

## 3 Qualitative Analysis of the Model

We characterize the existence and stability of equilibria of the system described by equations (5a)–(5e).

### 3.1 Equilibrium Analysis

At equilibrium, all time derivatives vanish. The antibiotic equations (5d) are decoupled and yield the unique solutions 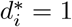 for *i* = 1, 2, 3. Substituting these values into the remaining equations and setting the derivatives to zero gives the algebraic system:

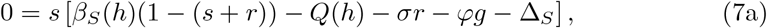

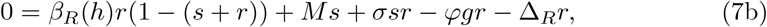

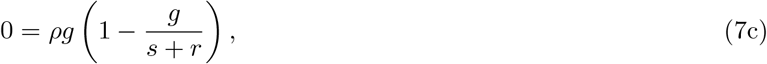

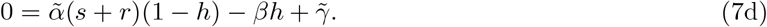

For analytical tractability, the dietary perturbation, we replace 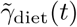 by its temporal average 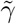. Equation (7c) yields two distinct biological scenarios: the absence of immune response (*g*^∗^ = 0) or an active immune response proportional to the total bacterial load (*g*^∗^ = *s*^∗^ + *r*^∗^). Equation (7d) can be solved explicitly for the equilibrium pH:

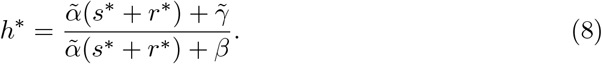

Equation 8 shows that the equilibrium pH increases monotonically with the total bacterial population (*s*^∗^ + *r*^∗^), reflecting the cumulative buffering effect of bacterial urease activity.

We proceed the analysis by examining the two immune response cases.

#### 3.1.1 Case 1: Absent Immune Response

With *g*^∗^ = 0, the system reduces to

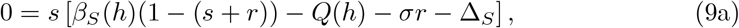

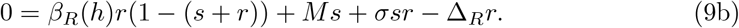

We identify three possible equilibria.

#### Infection-Free Equilibrium (IFE)

The trivial solution *s*^∗^ = 0, *r*^∗^ = 0 exists for all parameter values. From (8), the corresponding pH is 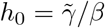. Thus,

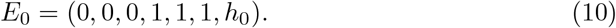

#### Resistant-only equilibrium

Setting *s*^∗^ = 0 and *r*^∗^ *>* 0 in (9b) gives the condition *β*_*R*_(*h*)(1 −*r*^∗^) = Δ_R_ that implies 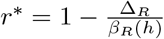 that implies *β*_*R*_(*h*)*/*Δ_R_ *>* 1 leading to the following definition

##### Definition 1

(Fitness of Resistant Bacteria) *The basic reproductive number for the resistant bacterial population in the absence of immune response and sensitive bacteria is:*

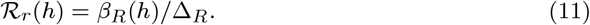

*It represents the average number of new resistant bacteria produced by a single resistant bacterium during its lifespan*.

Thus, *r*_1_ = 1 − 1*/*ℛ_*r*_(*h*_1_) = (ℛ_*r*_(*h*) − 1)*/* ℛ_*r*_(*h*) and the equilibrium is:

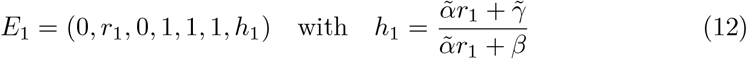

that exists if and only if ℛ_*r*_(*h*_1_) *>* 1, indicating that the resistant strain can sustain itself in the absence of competition and immune pressure.

#### Coexistence equilibrium

For *s*^∗^ *>* 0 and *r*^∗^ *>* 0, Equaton (9a) can be rearrange as:

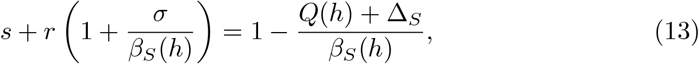

showing that bacteria coexistence required *β*_*S*_(*h*)*/*(*Q*(*h*) + Δ_S_) *>* 1 inducing the following definition:

##### Definition 2

(Fitness of Sensitive Bacteria) *The effective reproductive number for the sensitive bacterial population, accounting for antibiotic pressure and mutation, is:*

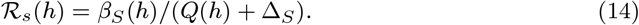

*It represents the average number of new sensitive bacteria produced by one sensitive bacterium that survives all elimination pressures except competition from resistant bacteria*.

Existence of sensitive bacteria therefore requires ℛ_*s*_(*h*) *>* 1. From Equation (13):

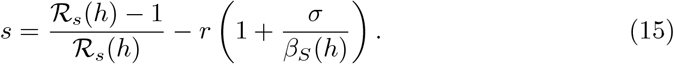

and substituting into (9b) yields the quadratic equation in *r*

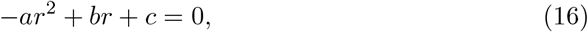

where the coefficients are: 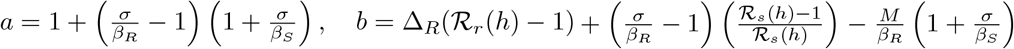 and 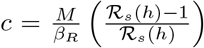. The coefficient *a* simplifies to a positive quantity: 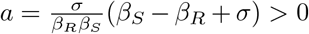, as *β*_*S*_ ≥ *β*_*R*_ (fitness cost of resistance) and all parameters are positive. The term *c* is positive if ℛ _*s*_(*h*) *>* 1 and the quadratic (16) has a positive discriminant and therefore a unique positive root *r*^∗^. Substituting *r*^∗^ back into Equation (15) gives *s*^∗^. Important to note that *s*^∗^ *>* 0 requires 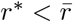, where 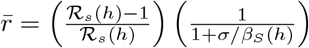. The existence of equilibrium

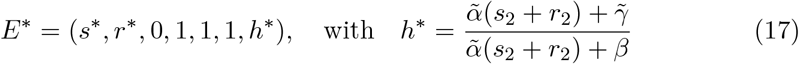

further requires that the resistant population is not overly fit, formalized by a threshold derived from the condition 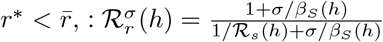 . such that *E*^∗^ exists if ℛ_*s*_(*h*^∗^) *>* 1 and 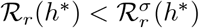.

#### 3.1.2 Case 2: Active Immune Response

With an active immune response (*g*^∗^ = *s*^∗^ + *r*^∗^), the system becomes:

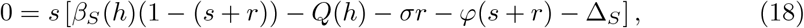

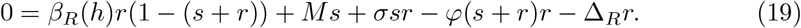

#### Resistant-only equilibrium with immunity

Assuming *s*^∗^ = 0, equation (19) simplifies to

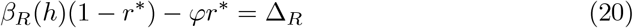

Solving for *r*^∗^ gives: 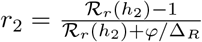. The equilibrium

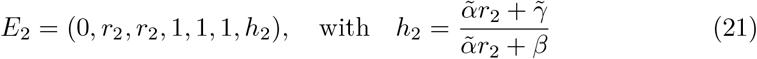

exists if and only if ℛ_*r*_(*h*_2_) *>* 1. The immune response reduces the equilibrium bacterial load compared to *E*_1_.

#### Coexistence Equilibrium with Immunity

For *s*^∗^ *>* 0 and *r*^∗^ *>* 0, we obtain from (18) a relationship analogous to (15) but modified by the immune clearance term *φ*.

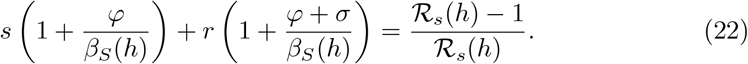

Again, ℛ_*s*_(*h*) *>* 1 is required. Substituting into (19) yields another quadratic equation in *r* with a unique positive root *r*_3_.

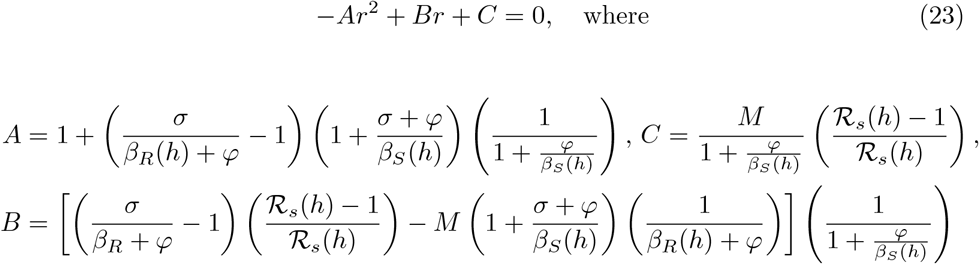

*A* can be simplified using basis algebra as 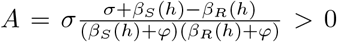 and under the condition ℛ_*s*_(*h*) *>* 1, *C >* 0, the discriminant of the quadratic equation (23) is positive and therefore, it has a unique positive root *r*_3_

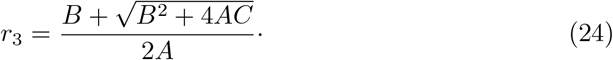

Substituting back *r*_3_ into equation 22 gives *s*_3_ and then *g*_3_ = *r*_3_ + *s*_3_. The equilibrium

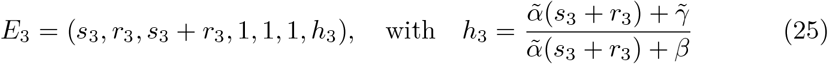

existence is governed by a second immune-related threshold 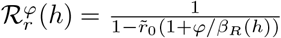 with 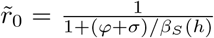, such that *E*_3_ exists if 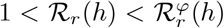 or if ℛ_*r*_(*h*) *>* 1, but 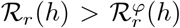 when 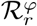 is negative or if ℛ_*r*_(*h*) ≤ 1. Biologically this equilibrium represents a state of chronic, balanced polymicrobial infection under immune surveillance.

The following theorem summarizes the different equilibrium solutions and their existence criterion.

##### Theorem 1

(Existence of Equilibria) *The in-host model has the following equilibria in the domain* Ω:

1. ***Infection-free equilibrium*** *E*_0_: (0, 0, 0, 1, 1, 1, *h*_0_). *Always exists*.
2. ***Resistant-only equilibrium*** *E*_1_: (0, *r*_1_, 0, 1, 1, 1, *h*_1_). *Exists iff ℛ*_*r*_(*h*_1_) *>* 1.
3. ***Coexistence equilibrium (No Immunity)*** *E*^∗^: (*s*^∗^, *r*^∗^, 0, 1, 1, 1, *h*^∗^). *Exists iff ℛ*_*s*_(*h*^∗^) *>* 1 *and* 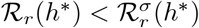.
4. ***Resistant-only equilibrium (With Immunity)*** *E*_2_: (0, *r*_2_, *r*_2_, 1, 1, 1, *h*_2_). *Exists iff ℛ*_*r*_(*h*_2_) *>* 1.
5. ***Coexistence equilibrium (With Immunity)*** *E*_3_: (*s*_3_, *r*_3_, *s*_3_ + *r*_3_, 1, 1, 1, *h*_3_). *Exists iff* ℛ_*s*_(*h*_3_) *>* 1 *and*
  i. 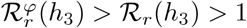, *or*
  ii. *ℛ*_*r*_(*h*_3_) *>* 1, *but* 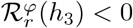, *or*
  iii. *ℛ*_*r*_(*h*_3_) ≤ 1.

#### 3.1.3 Biological Interpretation of Thresholds

The analysis reveals that the system’s long-term behavior is governed by two pH-dependent reproductive numbers: ℛ_*s*_(*h*) for sensitive bacteria and ℛ_*r*_(*h*) for resistant bacteria. Successful eradication (convergence to *E*_0_) requires therapeutic conditions that drive both ℛ_*s*_(*h*) *<* 1 and ℛ_*r*_(*h*) *<* 1. If ℛ_*r*_(*h*) *>* 1 but ℛ_*s*_(*h*) *<* 1, treatment fails and the resistant strain dominates (*E*_1_ or *E*_2_). If both exceed unity, coexistence (*E*^∗^ or *E*_3_) is possible, with the immune response determining the specific steady state. The thresholds 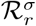 and 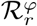 delineate the competitive balance between the strains under different immune contexts.

### 3.2 Local Stability Analysis

The local stability of the equilibria identified in Section 3.1 is determined by analyzing the eigenvalues of the system’s Jacobian matrix, evaluated at each equilibrium point. The dynamics of the antibiotic concentrations (*d*_1_, *d*_2_, *d*_3_) are decoupled and globally stable, with eigenvalues −*τ*_1_, −*τ*_2_, and −*τ*_3_. Therefore, we focus our stability analysis on the 4-dimensional subsystem (*s, r, g, h*). We denote the right-hand sides of equations (5a), (5b), (5c), and (5e) as *F*_1_, *F*_2_, *F*_3_, and *F*_4_, respectively. The Jacobian matrix *J* of this subsystem has thr general form:

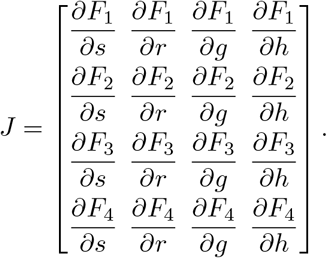

The explicit partial derivatives are provided in A.1. The sign of the real parts of the eigenvalues of *J*, evaluated at an equilibrium, determines its local asymptotic stability.

#### 3.2.1 Stability of the Infection-Free Equilibrium

At *E*_0_ = (0, 0, 0, *h*_0_), the Jacobian simplifies to a block-triangular form. Evaluating the derivatives with *g* = *s* + *r* yields:

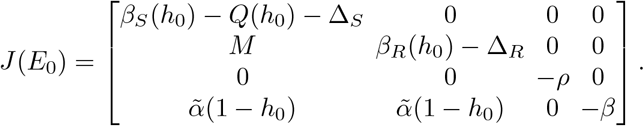

The eigenvalues of such matrix are the diagonal elements:

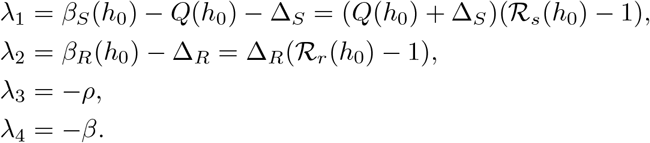

The following proposition characterizes the stability of the IFE state.

##### Proposition 2

(Stability of *E*_0_) *The infection-free equilibrium E*_0_ *is locally asymptotically stable within the biologically relevant subspace* (*s, r, g, h*) *if and only if* ℛ_*s*_(*h*_0_) *<* 1 *and* ℛ_*r*_(*h*_0_) *<* 1. *It is unstable if either* ℛ_*s*_(*h*_0_) *>* 1 *or* ℛ_*r*_(*h*_0_) *>* 1.

This condition biologically means that eradication is possible only when the reproductive capacity of both bacterial strains, under treatment pressure and at equilibrium pH *h*_0_, is less than one. Satisfying this requires that the combined effect of antibiotics (through *Q*(*h*_0_)) and natural clearance (Δ_S_) outweighs the replication capacity of sensitive bacteria, justifying the use of combination therapy to maximize *Q*(*h*_0_). Additionally, even if antibiotics are ineffective against resistant bacteria, through sufficient fitness cost and natural clearance. Successful treatment requires maintaining pH in a range that minimizes ℛ_*s*_(*h*) and ℛ_*r*_(*h*) while maximizing antibiotic efficacy.

#### 3.2.2 Stability of the Resistant-only Equilibrium without Immunity

The equilibrium *E*_1_ = (0, *r*_1_, 0, *h*_1_) exists when ℛ_*r*_(*h*_1_) *>* 1. Its Jacobian matrix takes the form:

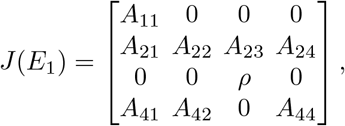

where by using equilibrium condition *β*_*R*_(*h*_1_)(1 − *r*_1_) = Δ_R_,

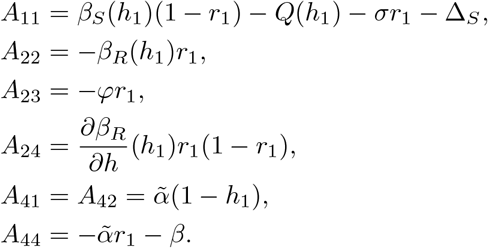

The eigenvalues are *λ*_1_ = *A*_11_, *λ*_2_ = *A*_22_, *λ*_3_ = *ρ*, and *λ*_4_ = *A*_44_. We have *λ*_2_ *<* 0, *λ*_4_ *<* 0, and *λ*_3_ = *ρ >* 0. The positive eigenvalue *λ*_3_ indicates that any infinitesimal immune activation will grow, driving the system away from *E*_1_. Therefore, *E*_1_ is unstable in the full system. This aligns with the biological expectation that a persistent bacterial load will eventually activate an adaptive immune response.

Hence the following theorem:

##### Theorem 3

(Stability of Resistant-only equilibrium *E*_1_) *The resistant-only equilibrium without immune response E*_1_ *is always unstable in the biological domain* Ω.

#### 3.2.3 Stability of the Coexistence Equilibrium without Immunity

A similar argument applies to *E*^∗^ = (*s*^∗^, *r*^∗^, 0, *h*^∗^). The Jacobian at *E*^∗^ always has an eigenvalue *λ* = *ρ >* 0 (see Appendix A.2) corresponding to the immune response direction. Consequently, we have the following result:

##### Proposition 4

*The coexistence equilibrium without an immune response, E*^∗^, *is always unstable in the full system that includes the immune response dynamics, due to the positive eigenvalue λ* = *ρ*.

The equilibria *E*_1_ and *E*^∗^ may, however, be relevant as transient states or as attractors in a hypothetical scenario of complete immune suppression.

#### 3.2.4 Stability of the resistant-only equilibrium with immunity

For *E*_2_ = (0, *r*_2_, *r*_2_, *h*_2_), which exists when ℛ_*r*_(*h*_2_) *>* 1, the Jacobian has a block structure due to *s* = 0:

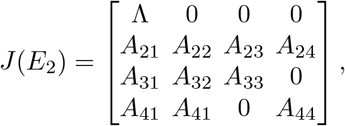

where Λ = *β*_*S*_(*h*_2_)(1 −*r*_2_) − *Q*(*h*_2_) −*σr*_2_ −*φr*_2 −_ Δ_S_, and the other entries are defined using the equilibrium conditions. One eigenvalue is simply *λ*_1_ = Λ. The remaining three eigenvalues are roots of the characteristic polynomial of the lower right 3 × 3 submatrix. The sign of *λ*_1_ = Λ determines the stability of *E*_2_ (see Appendix A.3) with respect to invasion by sensitive bacteria. Using the equilibrium condition for *r*_2_ (20), Λ can be expressed as:

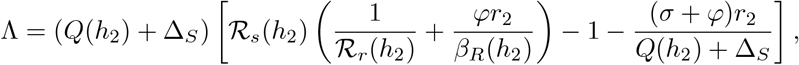

This leads to the following invasion condition:

##### Proposition 5

(Invasion condition for sensitive bacteria at *E*_2_) *The resistant-only equilibrium with immunity E*_2_ *is unstable to invasion by sensitive bacteria if* Λ *>* 0. *A sufficient condition for* Λ *>* 0 *(and thus instability) is* 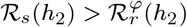, *where*

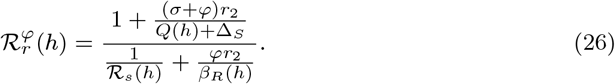

*Conversely, if* 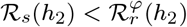, *then* Λ *<* 0 *and sensitive bacteria cannot invade*.

See the proof in Appendix A.4 .

For the remaining three eigenvalues, the Routh-Hurwitz criterion applied to the characteristic polynomial of the submatrix indicates that they have negative real parts for biologically plausible parameter values (see Appendix A.3). Therefore, the local stability of *E*_2_ hinges primarily on the sign of Λ.

#### 3.2.5 Stability of the coexistence equilibrium with immunity

The coexistence equilibrium *E*_3_ = (*s*_3_, *r*_3_, *s*_3_ + *r*_3_, *h*_3_) represents the most complex steady state. The 4 × 4 Jacobian matrix at *E*_3_ does not simplify into a block structure. Its characteristic polynomial is in the form:

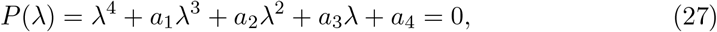

where the coefficients *a*_*i*_ are functions of the parameters and the equilibrium values. An analytical proof of stability for *E*_3_ is challenging due to the system’s complexity. However, our numerical simulations across a wide range of biologically plausible parameters consistently yield eigenvalues with negative real parts when exists (the Routh-Hurwitz criterion for Equation 27 is satified), suggesting that it is a locally asymptotically stable equilibrium.

#### 3.2.6 Summary of Stability Conditions

The local stability analysis yields the following key insights:

1. The clinically desirable infection-free state *E*_0_ is stable only when the treatment reduces the fitness of both bacterial strains below the critical threshold of one, i.e., ℛ_*s*_(*h*_0_) *<* 1 and ℛ_*r*_(*h*_0_) *<* 1.
2. Equilibria without an active immune response (*E*_1_ and *E*^∗^) are unstable in the presence of a recruitable immune system, as indicated by the positive eigenvalue *ρ*. to a modified threshold 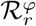 .
3. The resistant-only equilibrium with immunity *E*_2_ is stable provided the sensitive strain cannot invade (Λ *<* 0). This occurs when the sensitive strain is less fit relative
4. The coexistence equilibrium *E*_3_, when it exists, is generally stable, representing a state of chronic, balanced infection under immune surveillance.

This stability framework provides a theoretical foundation for understanding treatment outcomes. It predicts that therapeutic failure can manifest not only as the dominance of a resistant strain but also as a stable, mixed infection. The analytical conditions derived here will be validated and explored through numerical simulations in the following section.

## 4 Numerical Simulations and Results

To validate the analytical findings and explore the dynamical behavior of the pH-extended in-host model, we performed numerical simulations. This section details the numerical methodology, parameter selection, and presents results across four clinically relevant scenarios that illustrate the range of possible infection outcomes.

### 4.1 Numerical Method and Parameterization

The system of ordinary differential equations (5) was solved numerically using a fourth-order Runge–Kutta method with adaptive step size control, implemented in Python 3.9 (solve ivp function of SciPy). All simulations spanned 30 days to capture both transient dynamics and steady-state behavior. Initial conditions were set to represent an established infection: *s*(0) = 0.6, *r*(0) = 0.2, *g*(0) = 0.3, *d*_*i*_(0) = 0, and *h*(0) = 0.4, corresponding to a mixed bacterial population with moderate immune activation and slightly elevated gastric pH relative to baseline.

Parameter values were drawn from the literature where available (Table 1) and otherwise assumed with biological plausibility.

**Table 1.** Parameters of *H. pylori* infection model under antibiotic therapy, considering the immune response and the dynamic pH regulation. CFU stand for Colony-Forming Unit i.e a living H. pylori.

| Parameter | Definition | Value | Reference |
| --- | --- | --- | --- |
| $\beta_{S,\max}$ | Maximum growth rate of sensitive strain | 16.66 day<sup>-1</sup> | [9] |
| $\beta_{R,\max}$ | Maximum growth rate, resistant strain | 9.996 day<sup>-1</sup> | [9] |
| $\kappa_{i0}$ | Maximum antibiotic killing rate | 36.0 day<sup>-1</sup> | [9] |
| $\nu_i$ | Half-saturation antibiotic concentration | $2.5 \times 10^{-4}$ mg mL<sup>-1</sup> | [9] |
| $\mu_1$ | Max. Mutation rate due to RIF effect | $6.6 \times 10^{-8}$ CFU<sup>-1</sup>day<sup>-1</sup> | [9] |
| $\mu_2$ | Max Mutation rate due to CIP effect | $3.8 \times 10^{-8}$ CFU<sup>-1</sup>day<sup>-1</sup> | [9] |
| $\mu_3$ | Max Mutation rate due to CLT effect | $3.0 \times 10^{-9}$ CFU<sup>-1</sup>day<sup>-1</sup> | [9] |
| $\eta$ | Horizontal gene transfer rate | $1.0 \times 10^{-5}$ CFU<sup>-1</sup>day<sup>-1</sup> | [9] |
| $\gamma$ | Immune clearance rate | $6.0 \times 10^{-6}$ cell<sup>-1</sup>day<sup>-1</sup> | [9] |
| $\delta_S$ | Natural death rates of sensitive strain | 0.0037 day<sup>-1</sup> | [9] |
| $\delta_R$ | Natural death rates resistant bacteria | 0.0037 day<sup>-1</sup> | [9] |
| $\zeta$ | Detachment rate | 0.5 day<sup>-1</sup> | [9] |
| $\rho$ | Immune recruitment rate (T-cells) | 0.6 day<sup>-1</sup> | [9] |
| $\phi$ | Lymphocyte recruitment capacity | 1.0 cell CFU<sup>-1</sup> | [9] |
| $\tau_1$ | Absorption rate (RIF) | 0.96 | [9] |
| $\tau_2$ | Absorption rate (CIP) | 0.45 | [9] |
| $\tau_3$ | Absorption rate (CLT) | 0.35 | [9] |
| $M_1$ | Maximum concentration (RIF) | $1.2 \times 10^{-3}$ mg mL<sup>-1</sup> | [9] |
| $M_2$ | Maximum concentration (CIP) | $2.5 \times 10^{-3}$ mg mL<sup>-1</sup> | [9] |
| $M_3$ | Maximum concentration (CLT) | 2.77 mg mL<sup>-1</sup> | [9] |
| $K$ | Bacteria carrying capacity | $2.1 \times 10^3$ CFU | [9] |
| *pH dynamic parameters* |  |  |  |
| $\alpha$ | Urease activity rate | 0.01 pH CFU<sup>-1</sup>day<sup>-1</sup> | Assumed |
| $\beta$ | Acid secretion feedback rate | 0.5 day<sup>-1</sup> | Assumed |
| $H_{\max}$ | Maximum urease pH achievable | 7.0 | [12] |
| $H_0$ | Baseline gastric pH | 2.50 | [12] |
| $H_{\text{opt}}$ | Optimal growth pH | 7.0 | [12] |
| $H_{\text{opt,abx}}$ | Optimal antibiotic pH | 5.5 | [16] |
| $\sigma_{\text{abx}}$ | Antibiotic pH tolerance | 1.0 | Assumed |
| $\lambda$ | Dietary decay rate | 2.0 day<sup>-1</sup> | Assumed |

A key feature is the pH-dependence of bacterial growth rates *β*_*S*_(*h*), *β*_*R*_(*h*) and antibiotic killing rates *κ*_*i*_(*h*), modeled as described in Section 2. The dietary perturbation 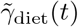 was modeled as three discrete meals per day, with pH perturbation strengths *γ*_1_ = 0.10, *γ*_2_ = 0.15, and *γ*_3_ = 0.08, corresponding to a light breakfast, a heavy lunch, and a very light supper, respectively. Each meal-induced pH change decays exponentially at rate *λ* = 2.0 day^−1^.

For each scenario, we verified the pH-dependent reproductive numbers ℛ_*s*_(*h*) and ℛ_*r*_(*h*) at the relevant equilibrium pH to confirm the analytical conditions for existence and stability. The thresholds 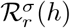 and 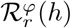 were computed accordingly.

### 4.2 Simulation Scenarios

We investigated four distinct scenarios, each characterized by specific combinations of the reproductive thresholds (Theorem 1) that dictate long-term outcomes:

1. **Scenario 1: Successful treatment and bacterial eradication**. Achieved by reducing the growth rate of resistant bacteria and increasing death rates (*β*_*R*,max_ = 0.664, *δ*_*S*_ = *δ*_*R*_ = 0.37). This corresponds to ℛ_*s*_(*h*) *<* 1 and ℛ_*r*_(*h*) *<* 1.
2. **Scenario 2: Treatment failure with resistant persistence**. Uses the baseline parameters in Table 1, yielding ℛ_*s*_(*h*) *<* 1 but ℛ_*r*_(*h*) *>* 1.
3. **Scenario 3: Bacterial coexistence under immunity**. Obtained by increasing both growth rates and reducing antibiotic efficacy (*β*_*S*,max_ = 19.0, *β*_*R*,max_ = 10.0, *κ*_*i*0_ = 0.9), leading to ℛ_*s*_(*h*) *>* 1 and ℛ_*r*_(*h*) *>* 1 while satisfying the coexistence condition 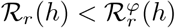.
4. **Scenario 4: Oscillatory dynamics under pH feedback**. A higher carrying capacity (*K* = 2.1 × 10^8^) perturbs the coexistence equilibrium. Compared to the base model [9], pH coupling reduces oscillation amplitude, introduces rapid damping, and confines residual oscillations to the resistant strain and immune response only.

### 4.3 Results and Interpretation

Figures 2–5 display the time series for each scenario. All four plots follow the same layout: sensitive bacteria, resistant bacteria, immune response, antibiotic concentrations, gastric pH, and dietary perturbation.

**Fig. 2.**
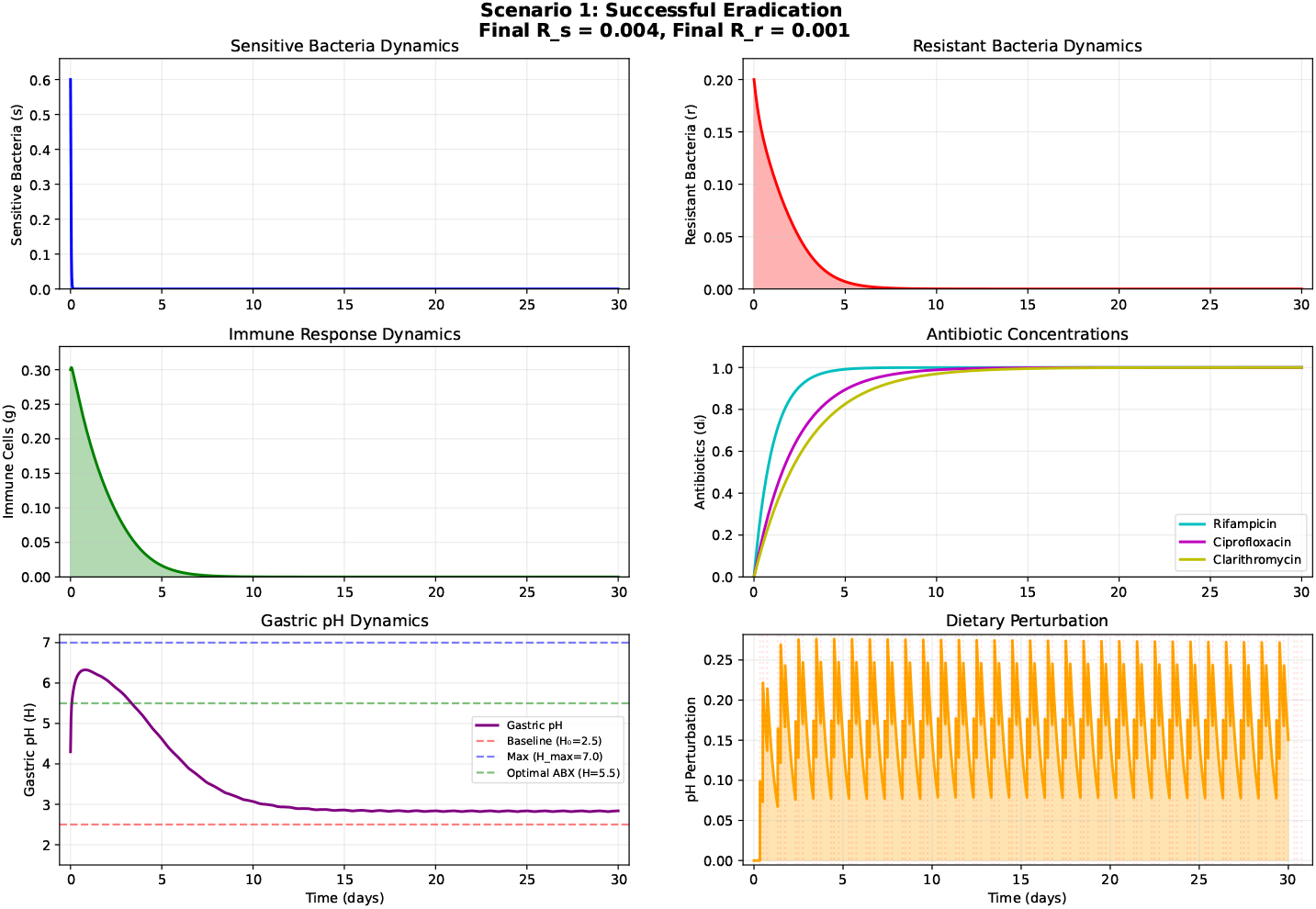
Scenario 1: Successful eradication. Both bacterial populations decline to zero within 10 days. Immune response peaks early and decays; gastric pH returns to baseline *H*_0_. Reproductive numbers at final state: *R*_*s*_ ≈ 0.004, *R*_*r*_ ≈ 0.001. This is characteristic of the convergence to the IFE (*E*_0_).

**Fig. 3.**
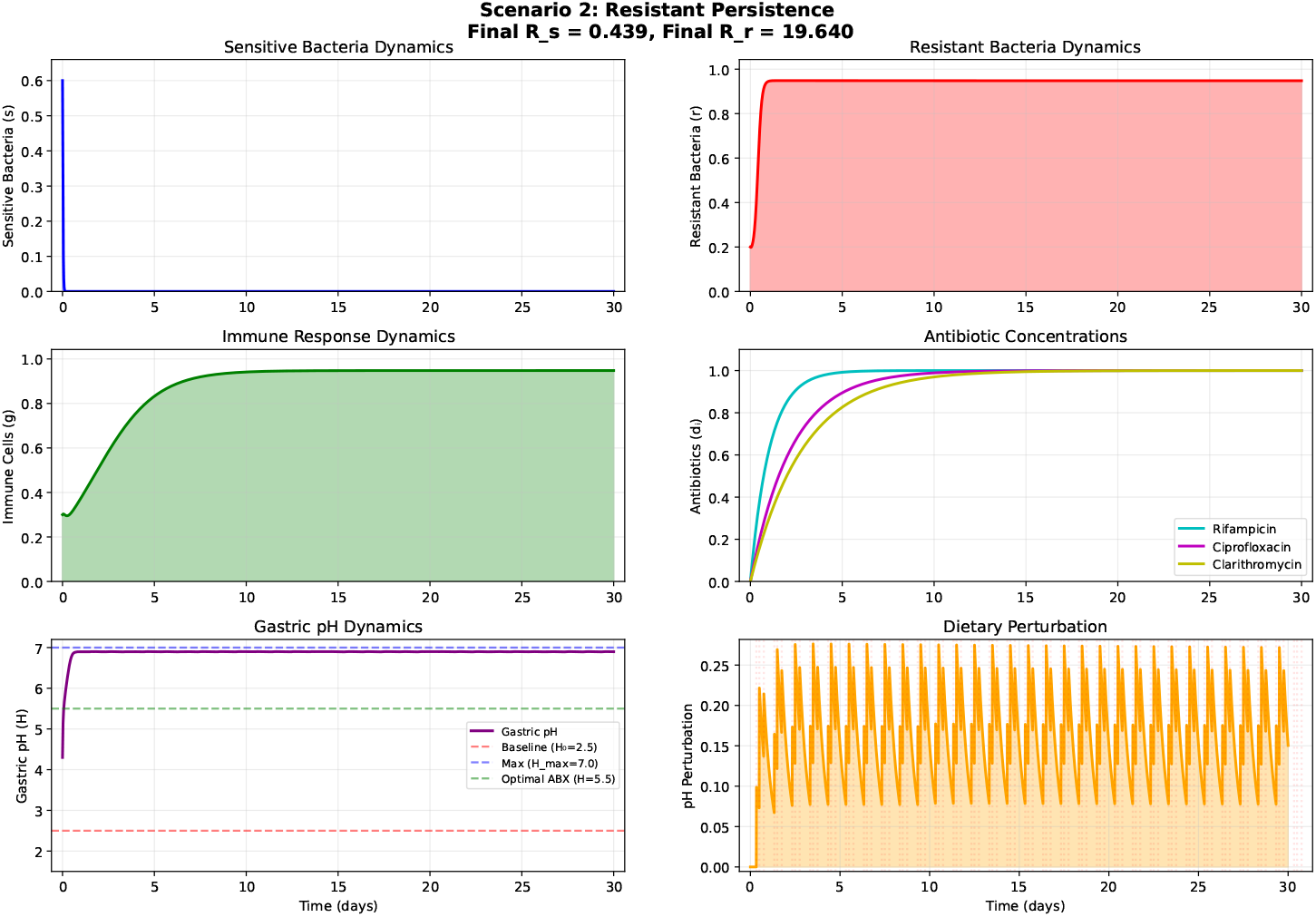
Scenario 2: Resistant persistence. Sensitive bacteria are eliminated, but resistant bacteria persist at a positive equilibrium (*r*_2_ ≈ 0.9). Immune response stabilizes at the same level; pH remains elevated (≈ 6.7). Final reproductive numbers: ℛ_*s*_ ≈ 0.44, ℛ_*r*_ = 19.64, 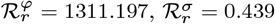.

**Fig. 4.**
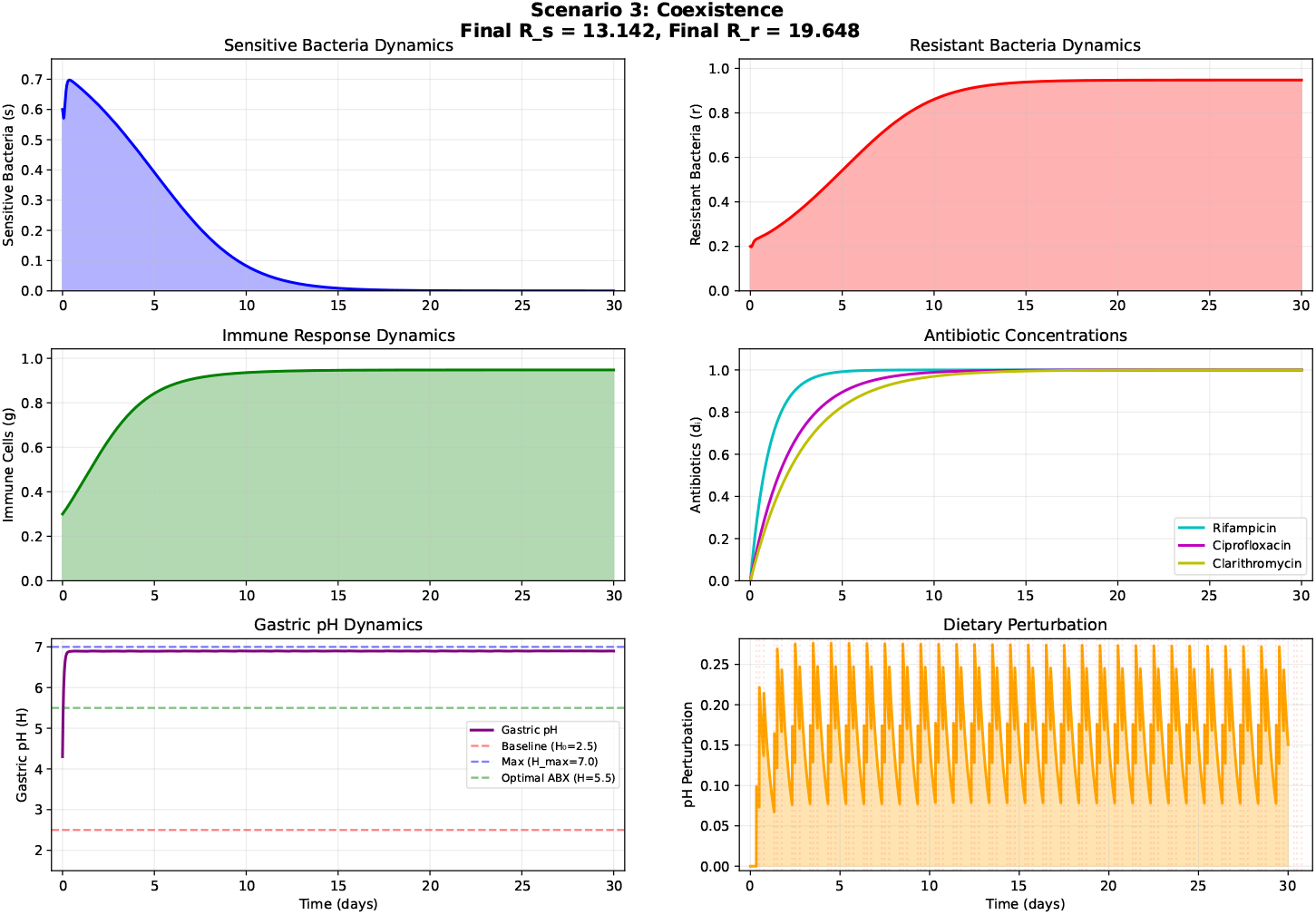
Scenario 3: Coexistence equilibrium *E*_3_. Both strains persist with immune response matching total bacterial load (*g* = *s* + *r*). pH stabilizes near neutrality. Thresholds: ℛ_*s*_ = 13.142, ℛ_*r*_ = 19.648, and 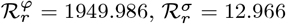 satisfying the coexistence condition.

**Fig. 5.**
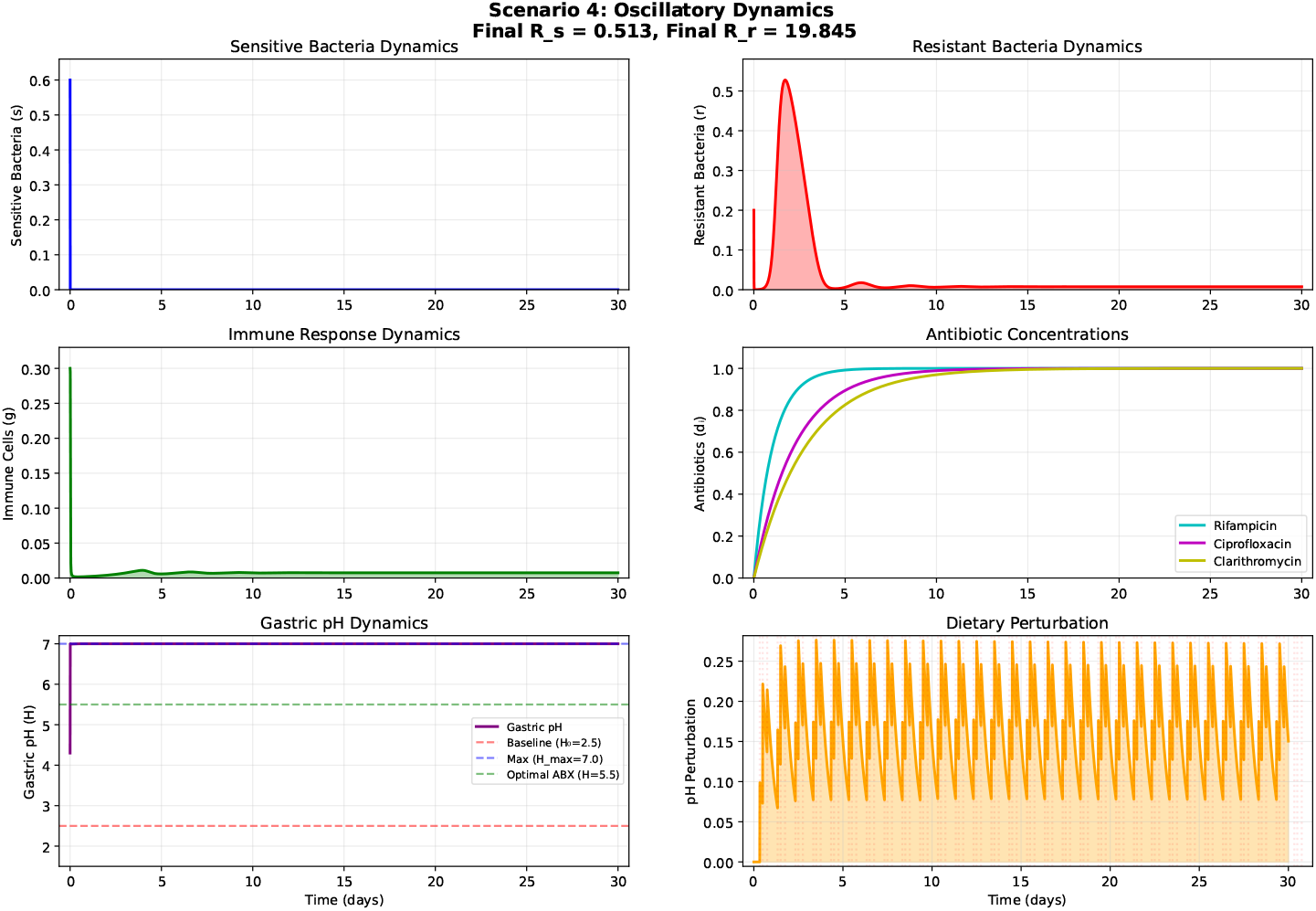
Scenario 4: Oscillatory dynamics under pH feedback. The system exhibits damped oscillations that transition rapidly to low-amplitude cycles, with residual oscillations confined to resistant bacteria and the immune response while pH stabilizes near neutrality. This contrasts with sustained oscillations in previous models [9] and suggests a mechanism for transient clinical relapse followed by stabilization. Final reproductive numbers: ℛ_*s*_ = 0.513, ℛ_*r*_ ≈ 19.845, 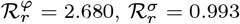.

In Scenario 1 (Fig. 2), both bacterial populations decline rapidly, confirming the stability of the infection-free equilibrium *E*_0_ when both reproductive numbers are below unity. The immune response transiently activates and then decays. Gastric pH returns to baseline (*h* →h_0_) as bacterial urease activity ceases. The antibiotics reach their maximum concentrations (*d*_*i*_ = 1) within the first few days.

Scenario 2 (Fig. 3) illustrates a common treatment failure mode: sensitive bacteria are cleared, but resistant bacteria survive and maintain an elevated pH, which further compromises antibiotic efficacy. The system converges to the resistant-only equilibrium *E*_2_, with immune response matching the resistant population (*g*→ *r*_2_).

Scenario 3 (Fig. 4) shows stable coexistence of both strains under immune surveillance. This polymicrobial state corresponds to equilibrium *E*_3_ and may represent a chronic infection that is difficult to eradicate. The pH remains high due to continued urease production by both strains.

Scenario 4 (Fig. 5) reveals a qualitatively different behavior: sustained oscillations with a period of approximately 15 days. These cycles arise from a Hopf bifurcation of the coexistence equilibrium *E*_3_ when the carrying capacity is sufficiently large. The interplay between bacterial growth, immune activation, and pH feedback generates recurrent peaks and troughs, potentially explaining relapsing–remitting clinical patterns.

### 4.4 pH-dependent fitness landscape

To further elucidate the role of gastric pH, we computed the reproductive numbers ℛ_*s*_(*H*) and ℛ_*r*_(*H*) across the physiological pH range for each scenario, together with the invasion thresholds 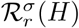 (plasmid transfer) and 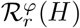 (immune-mediated). Figure 6 presents the graphs of these functions. The parameter values in the box on each figure correspond to the one used to simulate distinct clinical outcome above.

**Fig. 6.**
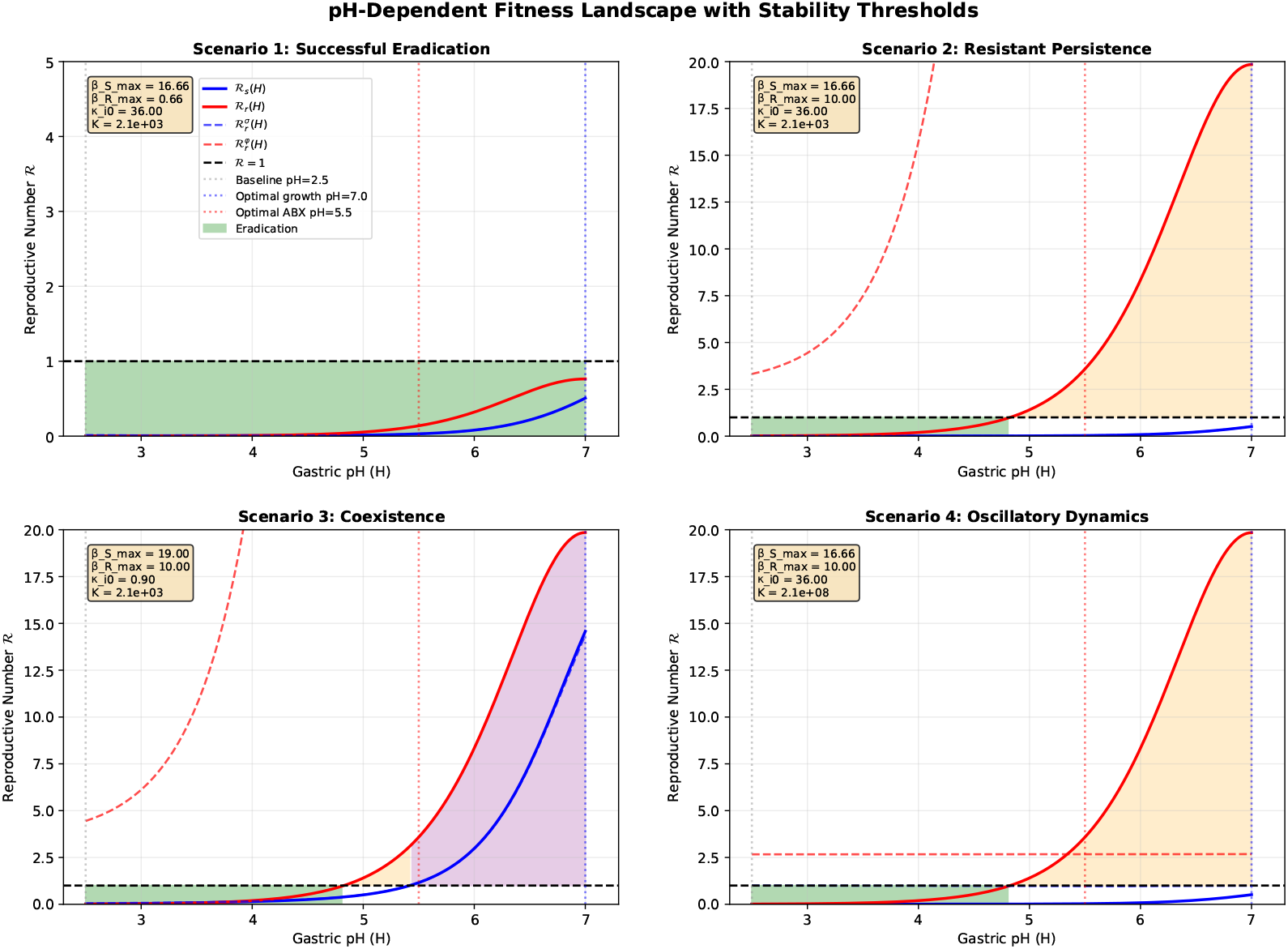
pH-dependent fitness landscape for the four scenarios. Colored zones indicate regions where different outcomes are predicted: green for eradication (ℛ_*s*_ *<* 1, ℛ_*r*_ *<* 1); orange for resistant persistence (ℛ_*s*_ *<* 1 *<* ℛ_*r*_ ); purple for coexistence (ℛ_*s*_ *>* 1, ℛ_*r*_ *>* 1); blue for sensitive persistence (rare). Dashed lines show the thresholds 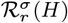 and 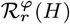. Vertical dotted lines mark baseline pH (*H*_0_ = 2.5), optimal growth pH (7), and optimal antibiotic pH (5.5). Parameter values for each scenario are displayed in the insets.

The landscape reveals distinct pH windows for each outcome. In Scenario 1, both reproductive numbers lie below unity across most of the pH range, with a narrow window of coexistence only near neutrality. Scenario 2 exhibits a wide region where ℛ_*r*_ *>* 1 while ℛ_*s*_ *<* 1, explaining the dominance of resistant bacteria. In Scenario 3, both ℛ _*s*_ and ℛ_*r*_ exceed unity for pH *>* 5, and the immune threshold 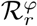 is sufficiently high to permit coexistence. Scenario 4 shows a similar structure but with the equilibrium destabilized, giving rise to oscillations.

The vertical lines highlight the tension between bacterial growth (optimal near pH 7) and antibiotic efficacy (optimal near pH 5.5). Successful therapy requires steering gastric pH into the eradication zone, a goal that may be achievable through a combination of acid suppression and timed antibiotic administration.

These numerical results confirm the analytical predictions and underscore the critical role of dynamic pH in shaping infection outcomes. The model not only reproduces expected clinical patterns but also reveals novel dynamical behavior (oscillations) that merits further experimental investigation.

## 5 Discussion

The in-host model presented in this study extends previous frameworks [9] by explicitly incorporating gastric pH as a dynamic state variable, coupling bacterial urease activity, host acid secretion, and diet changes. Our analysis reveals that pH is not a static background condition but an active ecological determinant that shapes the competitive landscape between antibiotic-sensitive and resistant *H. pylori* strains, modulates antibiotic efficacy, and ultimately dictates treatment outcomes. The numerical simulations confirm the analytical analysis and elucidate four distinct dynamical regimes, each with clear clinical correlates.

### 5.1 Interpretation of the four scenarios

**Successful eradication (Scenario 1)** demonstrates that when both pH-dependent reproductive numbers are driven below unity, the infection is cleared. This requires not only effective antibiotics but also a gastric pH that minimizes bacterial growth while maximizing drug activity. The rapid return of pH to baseline following bacterial clearance highlights the self-regulating nature of urease-driven alkalinization. This scenario underscores the rationale for combining acid-suppressive therapy with antibiotics: by raising pH toward the antibiotic optimum ( ≈5.5) while remaining below the growth optimum (7), anti-acid help satisfy the dual condition ℛ_*s*_ *<* 1 andℛ _*r*_ *<* 1.

**Resistant persistence (Scenario 2)** illustrates a common clinical failure mode: surviving resistant bacteria maintain elevated pH via urease activity, which compromises antibiotic efficacy and creates favorable growth conditions. This explains why incomplete eradication can lead to long-term resistant dominance.

**Coexistence (Scenario 3)** illustrates that both strains can persist over time under immune pressure, provided ℛ_*s*_ *>* 1 and ℛ_*r*_ *>* 1 and the immune-mediated threshold 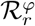 is satisfied. This polymicrobial state may represent chronic infections that are difficult to diagnose (since both strains are present) and may serve as reservoirs for horizontal gene transfer [17], potentially accelerating resistance spread. The model predicts that such coexistence is stable and robust to moderate perturbations, suggesting that once established, it may be challenging to disrupt without altering the pH landscape.

**Oscillatory dynamics (Scenario 4)** reveal how pH coupling modifies previously reported oscillations [9]. In our model, pH feedback reduces oscillation amplitude, induces rapid damping, and confines residual oscillations to the resistant strain and immune response only. When carrying capacity is sufficiently large, the equilibrium is perturbed by these modified oscillations, suggesting an instability consistent with a Hopf bifurcation. The oscillations, with a period of approximately 15 days, emerge from coupled feedback between bacterial growth, immune activation, and pH regulation. Clinically, such dynamics could manifest as relapsing-remitting symptoms, where patients experience periodic flare-ups followed by partial remission despite continuous antibiotic presence. Our model offers a testable hypothesis: pH elevation by urease activity creates a buffering effect that prevents uncontrolled proliferation, explaining why some patients experience transient symptoms that rapidly stabilize.

### 5.2 The pH-dependent fitness landscape

Figure 6 synthesizes the model’s predictions. The reproductive numbers ℛ_*s*_(*H*) and ℛ_*r*_(*H*) exhibit non-monotonic pH dependence, with growth optimal near neutrality and antibiotic efficacy optimal in the moderately acidic range. The intersection of these curves with the critical value ℛ= 1 defines pH windows for different outcomes. Several insights emerge:

1. **Therapeutic window:** There exists a narrow pH range (approximately 5.0–6.0) where ℛ_*s*_ *<* 1 and ℛ_*r*_ *<* 1 simultaneously, corresponding to successful eradication. This window lies between the growth optimum and the antibiotic optimum, explaining why moderate acid suppression (rather than complete neutralization) is beneficial.
2. **Thresholds matter:** The invasion thresholds 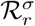 (plasmid transfer) and 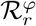 (immune-mediated) delineate the boundaries between coexistence and competitive exclusion. When 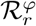 is low, even a moderately fit resistant strain cannot invade an established sensitive population; when it is high, coexistence is possible. This highlights the protective role of the immune system in preventing resistant dominance.
3. **pH as a control parameter:** Small changes in gastric pH can shift the system from one regime to another. For example, in Scenario 2, lowering pH from 6.5 to 5.5 would reduce ℛ_*r*_ below unity while keeping ℛ_*s*_ *<* 1, potentially converting failure into eradication. This suggests that precise pH modulation beyond simple acid suppression could be a therapeutic strategy. The diet can also therefore play a crucial role.

### 5.3 Dietary modulation of gastric pH

The pH-dependent fitness landscape (Fig. 6) reveals a critical threshold: when pH falls below approximately 4.7, both ℛ_*s*_(*H*) and ℛ_*r*_(*H*) drop below unity, indicating that sufficiently acidic conditions alone could suppress bacterial growth, regardless of antibiotic resistance status. This finding suggests a potential two-phase dietary strategy that warrants clinical investigation: (i) During initial treatment, interventions that elevate gastric pH into the therapeutic window (5.0–6.0) might optimize antibiotic stability and efficacy against sensitive strains. (ii) After treatment failure, When first-line therapy fails and resistant bacteria persist, maintaining a highly acidic gastric environment (pH *<* 4.7) could theoretically suppress the resistant population, potentially aiding eradication even without effective antibiotics. Several natural products have demonstrated urease-inhibiting properties in experimental studies [18], which may complement pH modulation by reducing the bacterium’s ability to neutralize gastric acid. Clinical trials have reported that certain dietary components can improve eradication success ([19], [18]), though the mechanisms remain incompletely understood and may involve multiple pathways beyond pH effects.

This dual strategy represents a hypothesis generated by our model: that extreme acidity (below 4.7) creates an inhospitable environment for *H. pylori*, while moderate acidity (5.0–6.0) enhances pharmacological killing. Dietary counseling as an adjunct to therapy remains speculative but merits prospective clinical investigation, particularly in resource-limited settings or after standard treatment failure.

### 5.4 Comparison with previous models

Our work builds directly on the model of Noguera et al. [9], which provided a robust framework for sensitive–resistant competition under antibiotic pressure and immune response. By adding dynamic pH, we have addressed a critical gap: the original model treated pH implicitly, assuming fixed antibiotic efficacy and growth rates. Our results show that pH feedback fundamentally alters the dynamics, particularly by damping oscillations and confining them to the resistant-immune loop.

Other modeling studies have considered pH in *H. pylori* dynamics [12, 20], but typically in a static or without under antibiotic therapy. To our knowledge, this is the first within-host model to couple pH mechanistically bacterial urease activity, host acid secretion, and dietary inputs under antibiotic therapy, and to derive pH-dependent reproductive numbers as explicit thresholds governing infection outcomes.

### 5.5 Clinical and therapeutic implications

Our findings have several practical implications for the clinical management of *H. pylori* infections. First, the model reinforces that combination therapy is essential. The condition for eradication, ℛ _*s*_ *<* 1 and ℛ_*r*_ *<* 1, requires both effective antibiotics to lower ℛ_*s*_ and either fitness costs or immune pressure to suppress resistant strains, meaning that monotherapy or inadequate regimens risk selecting for resistance.

Second, pH monitoring and modulation could play a direct role in guiding treatment. Routine measurement of gastric pH during therapy may help predict outcomes, as patients with pH consistently maintained within the eradication window (5.0–6.0) are likely to have higher success rates. The pH-dependent fitness landscape (Fig. 6) further reveals that maintaining pH below approximately 4.7 can directly suppress even resistant strains, a finding with immediate practical applications.

Several natural products have demonstrated urease-inhibiting properties in experimental studies [18], which may complement pH modulation by reducing bacterial acid adaptation. Clinical trials have reported that certain dietary components can improve eradication success [18, 19], though the mechanisms remain incompletely understood and may involve multiple pathways beyond pH effects. Dietary interventions, if validated, could offer affordable adjuncts to therapy, particularly in resource-limited settings.

The model’s oscillatory dynamics also suggest potential implications for treatment timing. In patients where oscillatory behavior occurs, administering antibiotics during the bacterial growth phase of the cycle might prove more effective than during the decline phase. a hypothesis that could be tested in controlled studies. More broadly, the feedback loop whereby resistant bacteria elevate pH and thereby protect themselves underscores the importance of resistance management. Early, aggressive therapy to prevent any resistant bacterial survival appears critical, as delayed or suboptimal treatment may inadvertently create conditions that favor resistant dominance.

Collectively, these implications position gastric pH as a central, modifiable variable in *H. pylori* management. The integration of pharmacological acid suppression, dietary pH modulation, and urease-inhibiting natural products offers a multi-pronged approach that could enhance eradication rates, particularly in the face of rising antibiotic resistance.

### 5.6 Limitations and future directions

Our model assumes a well-mixed gastric compartment, neglecting spatial pH gradients. Parameter uncertainty and the absence of stochastic effects also warrant caution.

Future extensions will address some of these limitations. We plan to incorporate spatial heterogeneity, and personalized diet to further refine therapeutic strategies against this resilient and globally prevalent pathogen. Additionally, we aim to explore the impact of different dosing schedules and combination regimens, using the pH-dependent reproductive numbers as a guide for optimization.

Despite these limitations, the current model provides a theoretically grounded and biologically plausible framework for understanding how gastric pH shapes *H. pylori* infection dynamics and treatment outcomes. It highlights pH not merely as a static barrier but as a dynamic, manipulable variable that sits at the center of host–pathogen–therapy interactions. The derivation of pH-dependent reproductive numbers ℛ_*s*_(*H*) and ℛ_*r*_(*H*) provides clear thresholds for bacterial persistence, while the model suggests that pH modulation, whether pharmacological or dietary could influence treatment outcomes. The hypothesis of a two-phase strategy (elevating pH during treatment to optimize antibiotic efficacy, and maintaining low pH after failure to suppress resistant strains) emerges from the model but requires rigorous clinical validation. By recognizing gastric pH as an active ecological force, we open new avenues for optimizing eradication strategies against this resilient and globally prevalent pathogen.

## Acknowledgements

The authors gratefully acknowledge the financial support of the African Union Commission through the Pan African University. We appreciate the support of Hiroshima University that hosted us during this research.

## Authors Contributions

**Alex Hermann Sockeng Koussok**: Conceptualization, Methodology, Analysis, Simulation, Writing–original draft. **Edward Richard Onyango**: Supervision, Methodology, Validation. **Koichi Fujimoto**: Supervision, Resources & Review. **Jean Jules Tewa**: Supervision, Project administration, reviewing.

## Declarations

### Data availability

This is a theoretical study and no empirical data were generated. All model assumptions and parameter values are fully described in the manuscript.

### Code availability

The Python code used to generate the simulation results and figures in this study is publicly available at: https://github.com/Hermann-Sockeng/H-pylori-model/releases/tag/v1.0.0. A permanent archived version with DOI is available at: DOI: 10.5281/zenodo.18739168

## Appendix A Supplementary: Mathematical Details

This appendix provides the complete mathematical details for the local stability analysis presented in Section 3.2.

### A.1 Jacobian Matrix of the biological-pH subsystem

The biological pH subsystem is defined by equations (5a), (5b), (5c), and (5e). Let *F*_1_, *F*_2_, *F*_3_, and *F*_4_ denote the right-hand sides of these equations, respectively. The Jacobian matrix *J* = [∂F_*i*_*/*∂x_*j*_], with **x** = (*s, r, g, h*) ^*T*^, is given by:

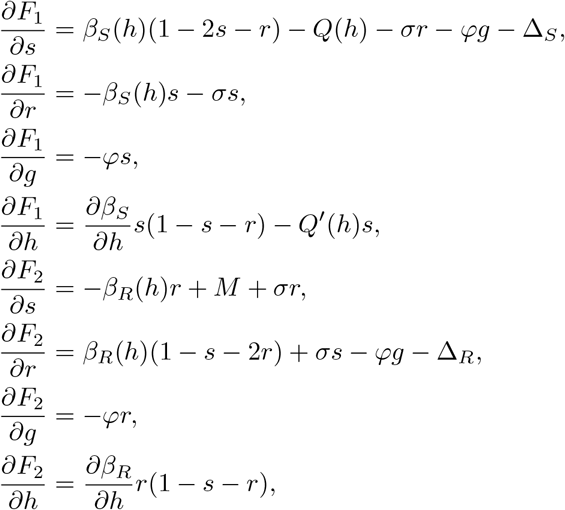

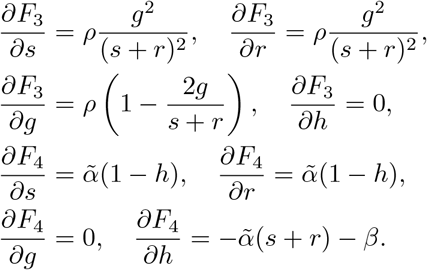

The pH-dependent functions are:

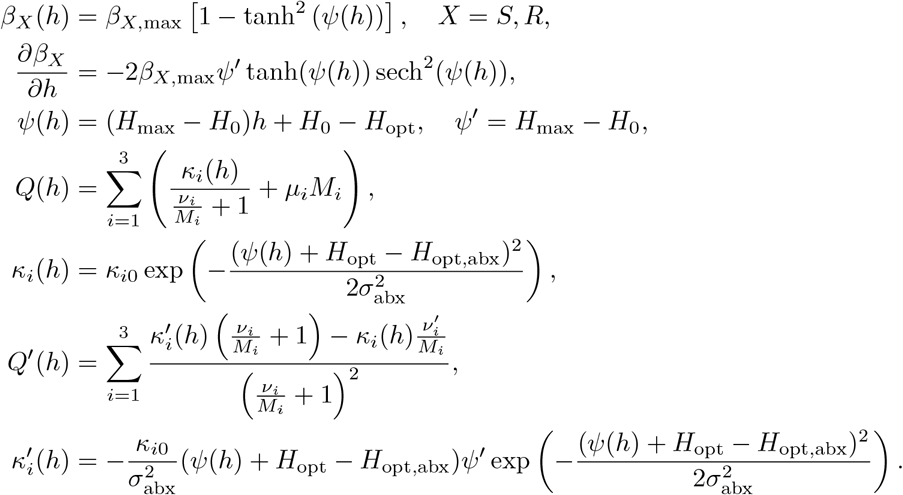

### A.2 Eigenvalue at the coexistence equilibrium without immunity

To obtain the eigenvalue *λ* = *ρ >* 0 for the equilibrium *E*^∗^ = (*s*^∗^, *r*^∗^, 0, *h*^∗^), we examine the structure of the Jacobian matrix *J*(*E*^∗^) of the biological-pH subsystem (*s, r, g, h*). The equilibrium *E*^∗^ is characterized by *g*^∗^ = 0 and *s*^∗^, *r*^∗^ *>* 0. The relevant partial derivatives at this point are:

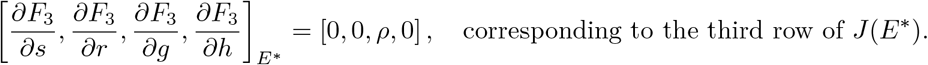

Now considering the full Jacobian evaluated at *E*^∗^:

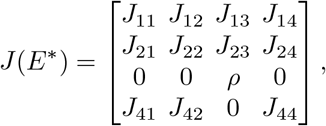

where *J*_*ij*_ represent the other partial derivatives in A.1 but not needed for this specific argument. The characteristic polynomial isgiven by:

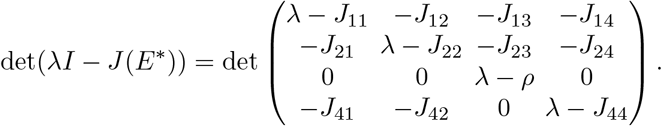

We can expand the determinant along the third row. The only non-zero entry in this row is the third element, (*λ* − *ρ*). The cofactor of this element is the determinant of the 3 *×* 3 matrix obtained by deleting the third row and third column:

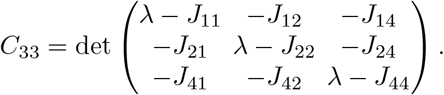

Therefore, the characteristic polynomial factors as:

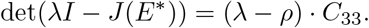

This factorization reveals that one eigenvalue of *J*(*E*^∗^) is simply *λ*_1_ = *ρ*.

### A.3 Eigenvalue Analysis for E2

For the equilibrium *E*_2_ = (0, *r*_2_, *r*_2_, *h*_2_), the Jacobian simplifies. Using the equilibrium condition *β*_*R*_(*h*_2_)(1 −*r*_2_) −*φr*_2_ = Δ_R_, the elements of the 3 × 3 submatrix for (*r, g, h*) are:

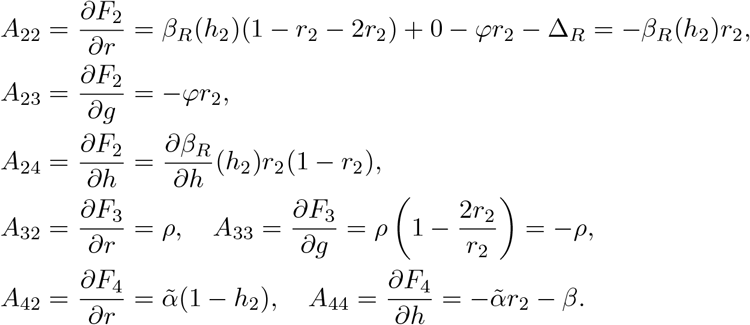

Thus, the submatrix is:

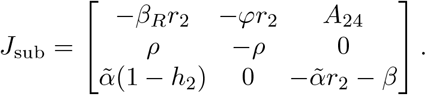

The characteristic polynomial of *J*_sub_ is:

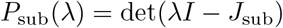

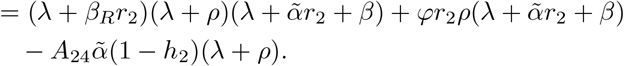

To apply the Routh-Hurwitz criterion, we examine the polynomial of the form

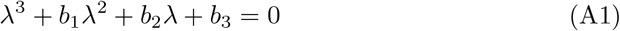

The coefficients are:

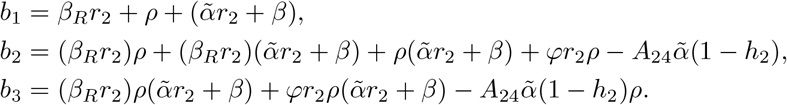

Our numerical simulation confirm that when *E*_2_ exists, the Routh-Hurwitz conditions for the cubic A1 is satisfy and these eigenvalues have negative real parts, and the stability is thus determined by the invasion eigenvalue 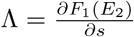.

### A.4 Proof of proposition 5

*Proof* The invasion eigenvalue at *E*_2_ is 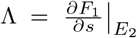. Using the expression for *J*_11_ and substituting the equilibrium conditions for *E*_2_ :

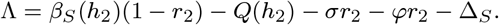

From the definition of ℛ_*s*_(*h*), we have *β*_*S*_ (*h*) = ℛ_*s*_(*h*)(*Q*(*h*) + Δ_S_ ). Substituting:

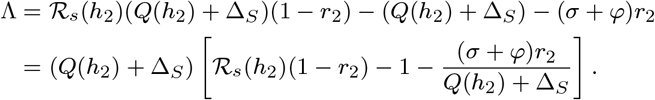

From *β*_*R*_(1 − *r*_2_) − *φr*_2_ = Δ_R_ (equilibrium condition 20) for r_2_, 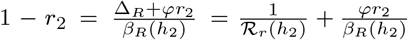 . Substituting this for (1 − *r*_2_):

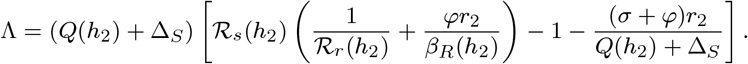

Then

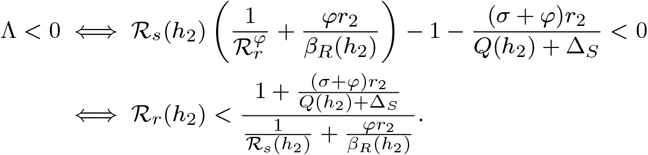

